# Temporal differentiation of resource capture and biomass accumulation as a driver of yield increase in intercropping

**DOI:** 10.1101/2021.02.17.431571

**Authors:** Nadine Engbersen, Rob W. Brooker, Laura Stefan, Björn Studer, Christian Schöb

## Abstract

- Intercropping, i.e. the simultaneous cultivation of different crops on the same field, has demonstrated yield advantages compared to monoculture cropping. These yield advantages have often been attributed to complementary resource use, but few studies quantified the temporal complementarity of nutrient acquisition and biomass production. Our understanding of how nutrient uptake rates of nitrogen (N) and phosphorous (P) and biomass accumulation change throughout the growing season and between different neighbors is limited.
- We conducted weekly destructive harvests to measure temporal trajectories of N and P uptake and biomass production in three crop species (oat, lupin and camelina) growing either as isolated single plants, in monocultures or as intercrops. Additionally, we quantified organic acid exudation in the rhizosphere and biological N_2_-fixation of lupin throughout the growing season. Logistic models were fitted to characterize nutrient acquisition and biomass accumulation trajectories.
- Nutrient uptake and biomass accumulation trajectories were curtailed by competitive interactions, resulting in earlier peak rates and lower total accumulated nutrients and biomass compared to cultivation as isolated single plants. Different pathways led to overyielding in the two mixtures. The oat–camelina mixture was characterized by a shift from belowground temporal niche partitioning of resource uptake to aboveground competition for light during the growing season. The oat–lupin mixture showed strong competitive interactions, where lupin eventually overyielded due to reliance on atmospheric N and stronger competitiveness for soil P.
- Synthesis: This study demonstrates temporal shifts to earlier peak rates of plants growing with neighbors compared to those growing alone, suggesting that the observed temporal shifts in our experiment are driven by competitive interactions rather than active plant behavior to reduce competition. The two differing pathways to overyielding in the two mixtures highlight the importance of examining temporal dynamics in intercropping systems to understand the underlying mechanisms of overyielding.

## 1. Introduction

Intercropping, i.e. the simultaneous growth of two or more species in the same field for all or part of their growing period, is a promising tool to sustainably maintain or increase yields by increasing diversity, thus maintaining natural ecosystem services and thereby limiting the input of agrochemicals (Lithourgidis *et al.* 2011; Brooker *et al.* 2016). Yield advantages in intercropping occur due to resource complementarity, where two or more species of an intercrop acquire different resources or acquire the same resources at different places belowground (Hauggaard-Nielsen, Ambus & Jensen 2001; Li *et al.* 2018) or at different times (Yu *et al.* 2015; Zhang *et al.* 2017; Dong *et al.* 2018). This reduces niche overlap and competition between individuals in the intercrop. To optimize intercropping systems, the aim is to select crops that differ in resource acquisition in time, space or form to maximize complementarity and reduce competition (Stomph *et al.* 2020).

Recent work has stressed the importance of including temporal dynamics of plant–plant interactions into competition studies (Trinder *et al.* 2012; Zhang *et al.* 2017). While most studies examining the mechanisms underlying dynamic plant–plant interactions have focused on temporal segregation of biomass accumulation, some also include measurements of essential nutrient uptake rates. For instance, Zhang *et al.* (2017) observed temporal niche differentiation between peak N uptake rates in a wheat/barley–maize intercropping system, where maximum N uptake rates of maize when intercropped with wheat/barley were delayed compared to when sole cropped. However, as maize growth was impaired during the co-growth period with wheat/barley (Zhang *et al.* 2015), the observed temporal shift towards later N uptake rates of intercropped maize is likely to be a passive response to early suppression rather than maize actively changing its N uptake rate to minimize interspecific competition. To differentiate between active and passive responses of plants to neighbors, it is crucial to associate temporal shifts of nutrient uptake rates with changes in plant productivity. More specifically, only when temporal shifts of nutrient uptake of a species are associated with a corresponding increase in productivity, there is evidence for active plant behavior towards niche complementarity. In contrast, temporal shifts in resource uptake associated with reduced productivity are most likely the sheer consequence of competitive suppression. Thus, studies of temporal dynamism of nutrient uptake and biomass in intercrops – although to date rare – are extremely valuable in helping us discover plant behavior for niche complementarity and potential mechanisms for yield benefits of intercropping.

Cereal–legume intercrops are usually very effective combinations in intercropping due to complementary N uptake strategies or facilitation in P uptake. The cereal is usually the stronger competitor for soil N and forces the legume neighbor to rely more heavily on atmospherically fixed N_2_ (Hauggaard-Nielsen *et al.* 2009), resulting in complementary N use. Moreover, P facilitation has been observed in several cereal–legume intercropping systems such as wheat–lupin and wheat–chickpea, where lupin and chickpea act as the P-mobilizing species (Li *et al.* 2003). By releasing carboxylates into the rhizosphere, the P-mobilizing species can increase the availability of inorganic P in the soil for itself but also for neighboring plant species (Li *et al.* 2014), thereby facilitating P uptake of the neighbor. Hence, complementary processes of both N and P capture are also involved in increasing productivity of cereal–legume intercrops. However, to what extent temporal dynamics in resource acquisition contribute to the success of cereal–legume mixtures is poorly understood. Furthermore, little is known about species mixtures of cereals with non-legumes, particularly cereals intercropped with crop species of the Brassicaceae family. The advantages of intercropping with Brassicaceae species can be multifaceted as Brassicaceae are known to show allelopathic activity, which can potentially be utilized to limit weed pressure, manage crop pests and diseases or even promote crop growth (Rehman *et al.* 2018).

The objectives of this study were to map trajectories of nutrient uptake and crop growth in two different intercropping systems to assess whether i) temporal differentiation in resource uptake and biomass accumulation can explain yield benefits in intercropping systems, ii) temporal differentiation in resource acquisition was reinforced through adjustments of the uptake pattern of species depending on neighbor identity, and iii) intra-specific adjustments in temporal nutrient uptake patterns contribute to yield benefits. Understanding these dynamics will help to improve our ability to predict successful species combinations for intercropping systems and to maximize the advantages of intercropping as a realistic alternative to current monoculture-based agricultural systems. To achieve this, we intercropped a cereal (oat (*Avena sativa*)) with either a legume (lupin (*Lupinus angustifolius*)) or a Brassicaceae (camelina (*Camelina sativa*)) and also – for comparison of temporal dynamics in inter- and monocrops – cultivated each crop species in a monoculture stand and as isolated single plants. For each species and planting pattern we examined intra- and inter-specific temporal differentiation by comparing maxima of nutrient uptake and biomass accumulation rates and assessed whether intra-specific temporal differentiation increased or decreased inter-specific temporal differentiation and yield benefits. Beyond nutrient uptake and biomass accumulation, we examined some particular physiological processes that could influence nutrient uptake patterns of intercropped species. Specifically, we investigated whether organic acids exudation by the legume or Brassicaceae differed temporally or quantitatively between crop species in intercrops, monocultures and isolated singles and whether it could be linked to increased P uptake of the exuding or neighboring crop species. We expected organic acids exudation of the legume or Brassicaceae to increase P uptake of the neighboring oat. Moreover, we traced biological N_2_-fixation of the lupin throughout the growing season to assess whether it differed between lupin intercropped with oat, in monoculture or as isolated single plant. Here, we expected biological N_2_-fixation to increase when lupin was intercropped with the cereal, due to increased competition for soil N by the cereal.

## 2. Materials and Methods

### 2.1. Site description

The study was carried out at the field site Aprisco de las Corchuelas, near Torrejón el Rubio, Cáceres, Spain. The site is located at 290 m a.s.l. (39°48’47.9” N 6°00’00.9” W). The regional climate is classified according to Köppen-Geiger (Kottek et al., 2006) as warm temperate, dry with hot summers. Total precipitation between February and June 2019 was 77.4 mm, daily average hours of sunshine during the growing season were 10.5 h and daily mean temperatures ranged between 9.6°C and 21.9°C, averaging 16°C. All climatic data are from the national meteorological service (www.aemet.es).

The experimental garden covered 120 m^2^, divided into 480 square plots of 0.25 m^2^ and 40 cm depth. Plots were open at the bottom and allowed root growth beyond 40 cm. The plots were arranged in 12 beds of 10 × 1 m, with two rows of 20 plots, resulting in 40 plots per bed. The plots were filled with local, unenriched agricultural soil. The soil consisted of 78% sand, 20% silt, 2% clay and contained 0.05% nitrogen, 0.5% carbon and 254 mg total P/kg with a mean pH of 6.3.

The experimental garden was irrigated throughout the growing season when plants required watering for survival. Irrigation was initiated when soil water content fell beneath 17% of field capacity at the rooting zone (10 cm below the soil surface). Irrigation events automatically increased soil water content to 25% of field capacity.

### 2.2. Experimental design

We used a complete randomized block design with three different crop species and three different diversity levels. The crop species were oat (*Avena sativa*), lupin (*Lupinus angustifolius*) and camelina (*Camelina sativa*) and the three diversity levels were monocultures, 2-species mixtures and isolated single plants. One block consisted of five plots: one plot of monoculture of each species, one plot with an oat–lupin mixture, one plot with an oat–camelina mixture. The isolated single plants of each species were arranged in a separate block and grown in a different bed to minimize neighbor effects. A monoculture plot consisted of four identical rows of the respective crop species. A mixture plot consisted of two alternating rows of each crop species. The sowing densities were: 400 seeds/m^2^ for oat, 160 seeds/m^2^ for lupin and 592 seeds/m^2^ for camelina and were based on current cultivation practice. Each block was repeated 54 times to allow for 18 destructive harvests with three replicates at each harvest. Sowing was done by hand on 2-3 February 2019.

### 2.3. Sample collection

After seedling emergence, a weekly destructive harvest took place, the first one on 21 February 2019 and the last one on 19 June 2019. Three individuals per plot and per species were randomly selected and marked. Roots of these three individuals were dug out carefully and gently shaken to remove soil. Root adhering soil (= rhizosphere soil (Veneklaas *et al.* 2003)) was gently brushed off and collected separately in 15 ml Falcon tubes. Afterwards roots were washed and stored in paper bags. Aboveground biomass of the three individuals was collected separately and separated into leaf, stem and – once available – fruits and seeds. All other individuals per species per plot were counted and aboveground biomass was harvested the same way as for the individuals and then pooled into one sample per species per plot. Aboveground biomass and roots were dried at 75 °C for at least 72h and weighed.

### 2.4. Nutrient analysis

Leaf biomass of the three individuals was pooled and ball milled to powder either in 1.2 ml tubes with two stainless steel beads in a bead mill (TissueLyserII, Qiagen) for three times 5 min or with a mixer mill (Mixer Mill MM 200, Retsch) for 30 seconds. Afterwards, either 100 mg (if available) or 4 mg (if the sample was too small) of ground leaf material was weighed into tin foil cups or 5 × 9 mm tin capsules and analyzed for N contents. The large samples (100 mg) were analyzed on a LECO CHN628C elemental analyzer (Leco Co., St. Joseph, USA) and the small (4 mg) samples on a PDZ Europa 20-20 isotope ratio mass spectrometer linked to a PDZ Europa ANCA-GSL elemental analyzer (Sercon Ltd., Cheshire, UK), respectively. All samples of lupin were analyzed on the mass spectrometer to obtain ^15^N data.

For P analysis, 100 mg of ground leaf material was weighed into microwave Teflon tubes and 2 ml H_2_O_2_ (30%) and 1 ml HNO_3_ (65%) were added. The samples were digested in a microwave (MLS-1200MEGA ETHOS) for ~25 min at a maximum temperature of 220 °C. The digests were diluted to a sample volume of 10 ml with Nanopure™ water and analyzed for P contents on an ICP-MS (Agilent 7900, Agilent Technologies, USA). For quality control we used the certified WEPAL (Wageningen Evaluating Programmes for Analytical Laboratories) reference materials IPE-100. Nutrient uptake was calculated as the product of nutrient concentration and aboveground biomass.

### 2.5. Organic acids analysis

The rhizosphere soil in the 15 ml Falcon tubes was immersed in 20 ml of a 0.2 mM CaCl_2_ solution and gently shaken for 30 min. The pH was measured in solution with a portable pH meter (Eutech pH 150, thermo). A subsample of solution was filtered into a 1.5 ml Eppendorf tube through a 0.2 μm syringe filter and acidified by adding a drop of 0.2 M H_2_SO_4_. All samples were kept frozen until analysis in Switzerland. Samples were analyzed on an IC (940 Professional IC Vario, Methrom) equipped with an ion exclusion column (PRP-X300 Ion Exclusion, Hamilton) and linked to an UV-VIS detector (UV-975, Jasco). The mobile phase was 0.5 mmol l^−1^ sulfuric acid with a flow rate of 2 ml min^−1^. The UV-VIS detector was connected to the IC with a 771 IC Compact Interface (Methrom) and the wavelength was set to 210 nm. Data processing was performed via MagIC Net Software (Methrom). Identification of organic acids was carried out by comparing retention time and absorption spectra with those of known standards.

### 2.6. Data analysis

All statistical analyses were performed in R version 3.6.0. (R Core Team 2019). To assess crop performance, we calculated the Land Equivalent Ratio (LER), defined as the sum of partial relative yields per species: 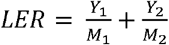, where Y_*i*_ is the yield of species *i* in the mixture, and M_*i*_ is the yield of species *i* in monoculture. Final seed yields from the final harvest week were used. LER values above 1 indicate a yield advantage of the mixture over the corresponding monocultures (Vandermeer 1989).

Temporal changes in biomass, and N and P accumulation were analyzed by fitting a logistic growth curve using non-linear least squares (nls) models (Trinder *et al.* 2012). Values of biomass, N and P from crop species in monoculture and mixture from weekly harvests were fitted to the following logistic model:

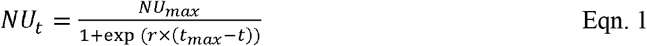

NU_t_ (biomass: g; P, N: mg) is the biomass accumulation or nutrient uptake of a crop species at the number of weeks after seedling emergence (t). NU_max_ (B_max_ for biomass) determines maximum cumulative nutrient uptake (biomass accumulation) of a crop species. r (day^−1^) is the relative nutrient uptake (biomass accumulation) rate. t_max_ is the time in weeks of reaching maximum nutrient (biomass) uptake rate. Starting values for the nls models were defined by first fitting a nls Levenberg-Marquardt model (nlsLM) and reusing the model parameters as starting values for the nls model. Eqn 1 was fitted separately to data for shoot biomass, N and P content across three replicates. Parameters indicating the fit of the models are reported in table S1 in the supplementary information.

The instantaneous biomass accumulation (g individual^−1^ week^−1^) or nutrient uptake (mg individual^−1^ week^−1^) can be derived as follows:

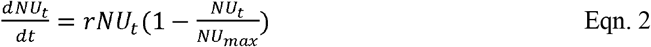

The maximum daily biomass accumulation and nutrient uptake rate (g day^−1^, mg day^−1^) which emerges at the time t_max_ was calculated as in Zhang *et al.* (2017) and is as follows: I_max_ = r × NU_max_/4.

Average ^15^N abundances from the reference plant (oat when available, otherwise camelina) at the same diversity level – i.e. oat from oat–camelina mixture as reference for mixtures, oat from oat monoculture as reference for monocultures and oat single plants as reference for single plants – and same harvest week were used to calculate the proportion of total aboveground lupin N derived from the atmosphere (% Ndfa) according to equation 3:

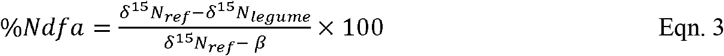

Where δ^15^N_ref_ was the average δ^15^N of the reference plant, δ^15^N_legume_ the average δ^15^N of the lupins and β was obtained in a separate greenhouse experiment. For this, single plants of *L. angustifolius* from the same seed material were grown in 5.5 l pots, filled with 0.7 – 1.2 mm coarse sand and inoculated with 100 ml soil suspension from the field site. Three replicates per harvest week were grown. Pots were watered twice daily with N-free McKnights solution (following the protocol of Unkovich et al. (2008)). Plants were harvested weekly and leaf samples were dried until constant weight, ground and analyzed for ^15^N as mentioned above. β values for each week are given in table S2.

## 3. Results

### 3.1. Crop performance

Mean LER values based on total plot-level grain yield of the final harvest week for the two mixtures were 1.84 ± 0.26 for oat–lupin and 1.63 ± 0.23 for oat–camelina, indicating that both mixtures overyielded. Partial LERs revealed that total plot-level grain yield of oat in both mixtures (oat– camelina: 1.47 ± 0.47, oat–lupin: 0.71 ± 0.05) and of lupin in the oat–lupin (1.12 + 0.4) mixture exceeded those of oat and lupin in monoculture, respectively. However, camelina in the oat–camelina (0.15 ± 0.08) mixture showed no yield benefits due to intercropping.

### 3.2. Intra-specific variability

No significant differences in biomass accumulation rates or nutrient uptake rates between mixtures, monocultures and isolated singles were observed for any species (Table 1), except for biomass accumulation rate of isolated single lupin, which was significantly higher than when the species was grown in a monoculture. However, isolated single plants always showed significantly higher maximum cumulative biomass (B_max_) and cumulative nutrient uptake (NU_max_) compared to the respective species grown in a community, except for N uptake of camelina (Table 1, Fig. S1).

**Table 1:**
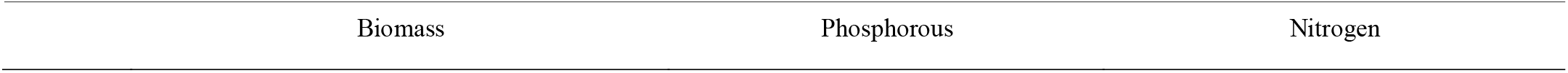
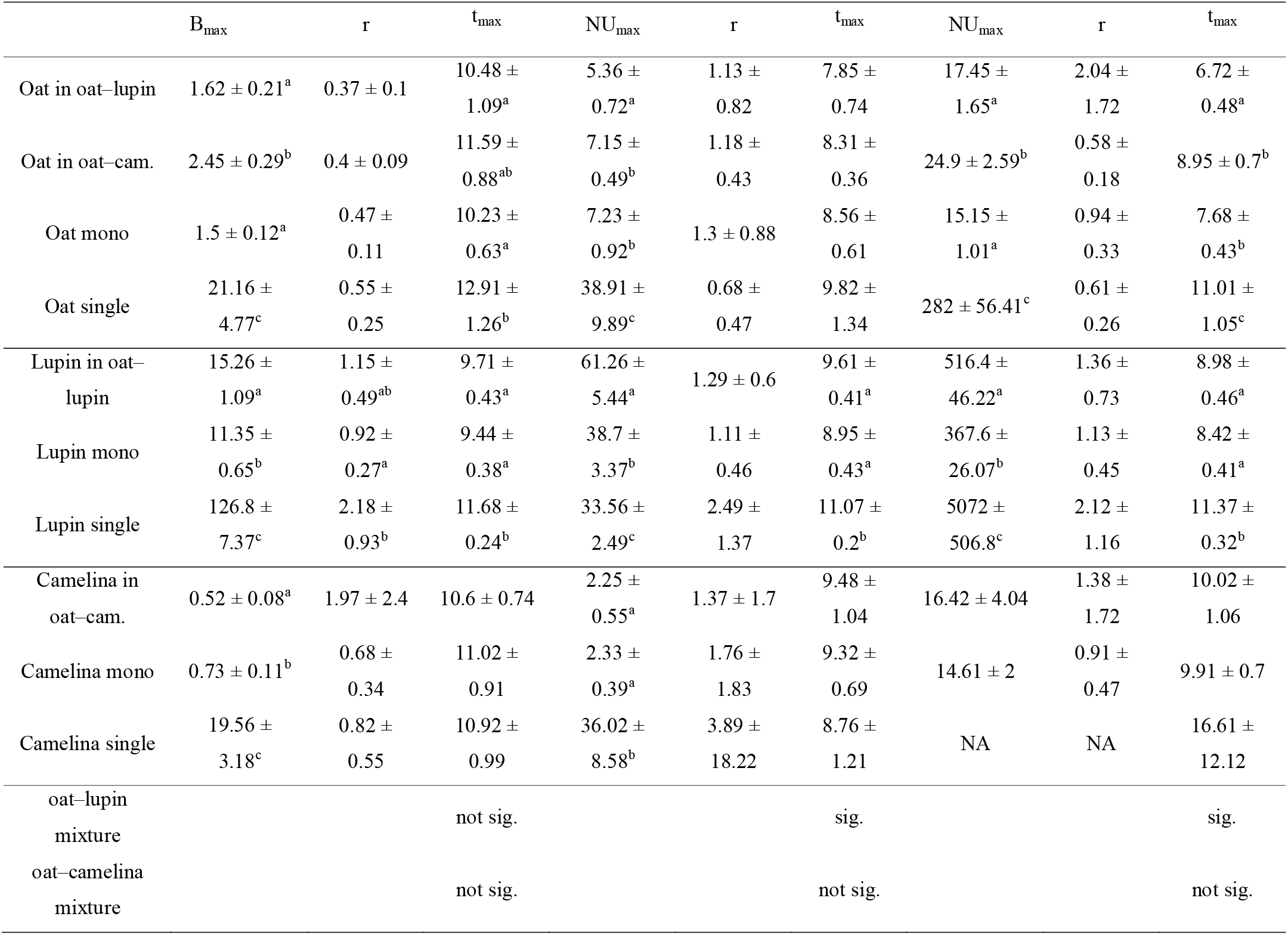
Mean values ± SE (n = 3) of model parameters fitted to logistic accumulation curves in Eqn. 1 using a nls model for biomass accumulation, N and P uptake. r is the rate constant of biomass production or nutrient capture (d^−1^), B_max_ is the maximum cumulative biomass (g), NU_max_ is the maximum cumulative nutrient uptake (mg) and t_max_ is the timepoint (number of weeks) at which the maximum biomass accumulation or maximum N or P uptake rate occurred. Parameter estimates with different superscripts within each subsetted column are significantly different between treatments. “Sig.” indicates significant differences in t_max_ between the two species intercropped in a mixture and “not sig.” indicates no significant differences. Significance is based on non-overlapping SE. Isolated singles of camelina were not fitted as they showed an exponential instead of logistic growth curve.

|  | B<sub>max</sub> | r | t<sub>max</sub> | NU<sub>max</sub> | r | t<sub>max</sub> | NU<sub>max</sub> | r | t<sub>max</sub> |
| --- | --- | --- | --- | --- | --- | --- | --- | --- | --- |
| Oat in oat–lupin | 1.62 ± 0.21<sup>a</sup> | 0.37 ± 0.1 | 10.48 ± 1.09<sup>a</sup> | 5.36 ± 0.72<sup>a</sup> | 1.13 ± 0.82 | 7.85 ± 0.74 | 17.45 ± 1.65<sup>a</sup> | 2.04 ± 1.72 | 6.72 ± 0.48<sup>a</sup> |
| Oat in oat–cam. | 2.45 ± 0.29<sup>b</sup> | 0.4 ± 0.09 | 11.59 ± 0.88<sup>ab</sup> | 7.15 ± 0.49<sup>b</sup> | 1.18 ± 0.43 | 8.31 ± 0.36 | 24.9 ± 2.59<sup>b</sup> | 0.58 ± 0.18 | 8.95 ± 0.7<sup>b</sup> |
| Oat mono | 1.5 ± 0.12<sup>a</sup> | 0.47 ± 0.11 | 10.23 ± 0.63<sup>a</sup> | 7.23 ± 0.92<sup>b</sup> | 1.3 ± 0.88 | 8.56 ± 0.61 | 15.15 ± 1.01<sup>a</sup> | 0.94 ± 0.33 | 7.68 ± 0.43<sup>b</sup> |
| Oat single | 21.16 ± 4.77<sup>c</sup> | 0.55 ± 0.25 | 12.91 ± 1.26<sup>b</sup> | 38.91 ± 9.89<sup>c</sup> | 0.68 ± 0.47 | 9.82 ± 1.34 | 282 ± 56.41<sup>c</sup> | 0.61 ± 0.26 | 11.01 ± 1.05<sup>c</sup> |
| Lupin in oat–lupin | 15.26 ± 1.09<sup>a</sup> | 1.15 ± 0.49<sup>ab</sup> | 9.71 ± 0.43<sup>a</sup> | 61.26 ± 5.44<sup>a</sup> | 1.29 ± 0.6 | 9.61 ± 0.41<sup>a</sup> | 516.4 ± 46.22<sup>a</sup> | 1.36 ± 0.73 | 8.98 ± 0.46<sup>a</sup> |
| Lupin mono | 11.35 ± 0.65<sup>b</sup> | 0.92 ± 0.27<sup>a</sup> | 9.44 ± 0.38<sup>a</sup> | 38.7 ± 3.37<sup>b</sup> | 1.11 ± 0.46 | 8.95 ± 0.43<sup>a</sup> | 367.6 ± 26.07<sup>b</sup> | 1.13 ± 0.45 | 8.42 ± 0.41<sup>a</sup> |
| Lupin single | 126.8 ± 7.37<sup>c</sup> | 2.18 ± 0.93<sup>b</sup> | 11.68 ± 0.24<sup>b</sup> | 33.56 ± 2.49<sup>c</sup> | 2.49 ± 1.37 | 11.07 ± 0.2<sup>b</sup> | 5072 ± 506.8<sup>c</sup> | 2.12 ± 1.16 | 11.37 ± 0.32<sup>b</sup> |
| Camelina in oat–cam. | 0.52 ± 0.08<sup>a</sup> | 1.97 ± 2.4 | 10.6 ± 0.74 | 2.25 ± 0.55<sup>a</sup> | 1.37 ± 1.7 | 9.48 ± 1.04 | 16.42 ± 4.04 | 1.38 ± 1.72 | 10.02 ± 1.06 |
| Camelina mono | 0.73 ± 0.11<sup>b</sup> | 0.68 ± 0.34 | 11.02 ± 0.91 | 2.33 ± 0.39<sup>a</sup> | 1.76 ± 1.83 | 9.32 ± 0.69 | 14.61 ± 2 | 0.91 ± 0.47 | 9.91 ± 0.7 |
| Camelina single | 19.56 ± 3.18<sup>c</sup> | 0.82 ± 0.55 | 10.92 ± 0.99 | 36.02 ± 8.58<sup>b</sup> | 3.89 ± 18.22 | 8.76 ± 1.21 | NA | NA | 16.61 ± 12.12 |
| oat–lupin mixture |  |  | not sig. |  |  | sig. |  |  | sig. |
| oat–camelina mixture |  |  | not sig. |  |  | not sig. |  |  | not sig. |

#### Oat

Maximum cumulative aboveground biomass of oat intercropped with camelina was 63% and 51% higher compared to oat in monoculture and oat intercropped with lupin, respectively (Fig. 1A, Table 1). Notably, despite biomass gains for oat grown with camelina, maximum cumulative P uptake was not different between oat in monoculture and oat intercropped with camelina but was 33% lower for oat intercropped with lupin (Fig. 1D). Cumulative P uptake of oat mixed with lupin was similar to that of the other two community combinations until week 8 and then increased more slowly than oat in monoculture and oat mixed with camelina (Fig. 1D). Cumulative N uptake of oat behaved similarly to biomass accumulation and showed 64% and 43% more accumulated N for oat when intercropped with camelina compared to oat monoculture and oat intercropped with lupin, respectively (Fig. 1G, Table 1). Between weeks 6 and 9, oat intercropped with lupin accumulated N at a faster rate than the other two community combinations, but accumulation came to a halt after week 9 while oat intercropped with camelina continued N uptake. The N uptake rate for oat in monoculture slowed after week 9 and came to a halt after week 11, resulting in the lowest final N accumulation of all combinations (Fig. 1G, Table 1).

**Fig. 1:**
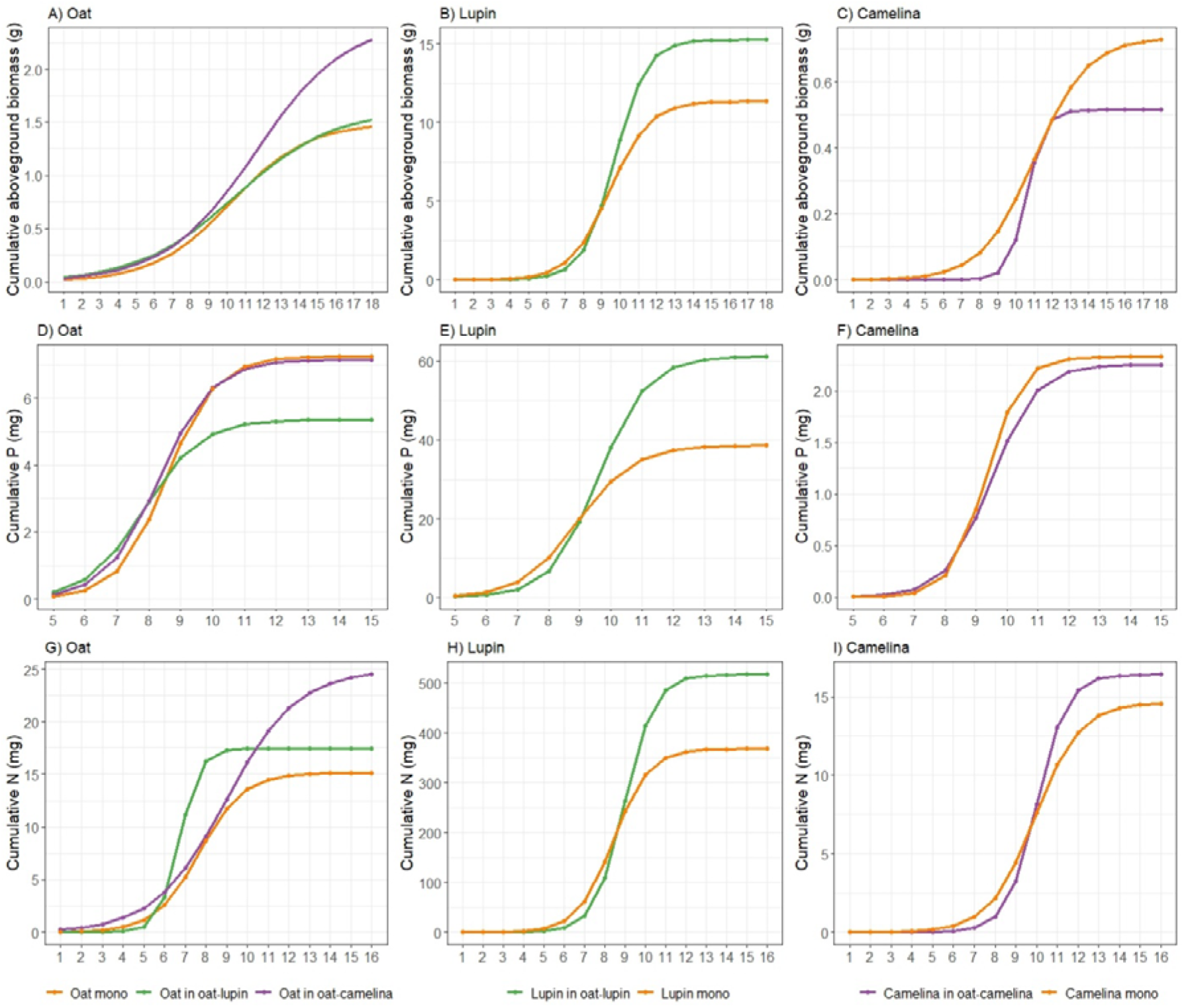
Trajectories of cumulative aboveground biomass (A-C), P (D-F) and N (G-I) uptake of oat with lupin (A, D, G: green), oat with camelina (A, D, G: purple), lupin (B, E, H) and camelina (C, F, I) when grown in mixture (purple, green) and in monoculture (orange). Curves were derived from Eqn. 1. Note different y-axis scales.

Maxima of instantaneous biomass accumulation and nutrient uptake (I_max_) of isolated single plants were considerably higher than in communities (Table S3). Maximum instantaneous biomass accumulation of oat grown in mixture with camelina exceeded the maxima of oat grown with lupin by 67% and oat in monoculture by 39% (Fig. 2A, B, Table S3). The timepoint of maximum instantaneous rates (t_max_) for biomass accumulation of isolated single oat occurred significantly later than of oat in monoculture and oat mixed with lupin but was not significantly different from oat mixed with camelina (Table 1). This indicated that, although not significantly, oat with camelina tended to have a later maximum instantaneous rate of biomass accumulation than oat in the other two community combinations (Fig. 2A, B, Table 1). Maximum instantaneous P uptake of oat mixed with camelina and in monoculture was, respectively, 40% and 55% higher than when intercropped with lupin (Table S3), while no differentiation between the timing of occurrence of maximum rates was observed (Fig. 2C, D, Table 1). Maximum instantaneous N uptake increased by 146 - 150% when oat was intercropped with lupin compared to when grown in monoculture or mixed with camelina (Fig. 2E, F, Table 1, S3). Timepoints of maximum instantaneous uptake rates of N in oat intercropped with lupin occurred one week earlier than when grown in monoculture and two weeks earlier than when intercropped with camelina. Timepoints of maximum instantaneous N uptake rates of isolated single oat occurred up to four weeks later than when oat was grown in a community (Table 1).

**Fig. 2:**
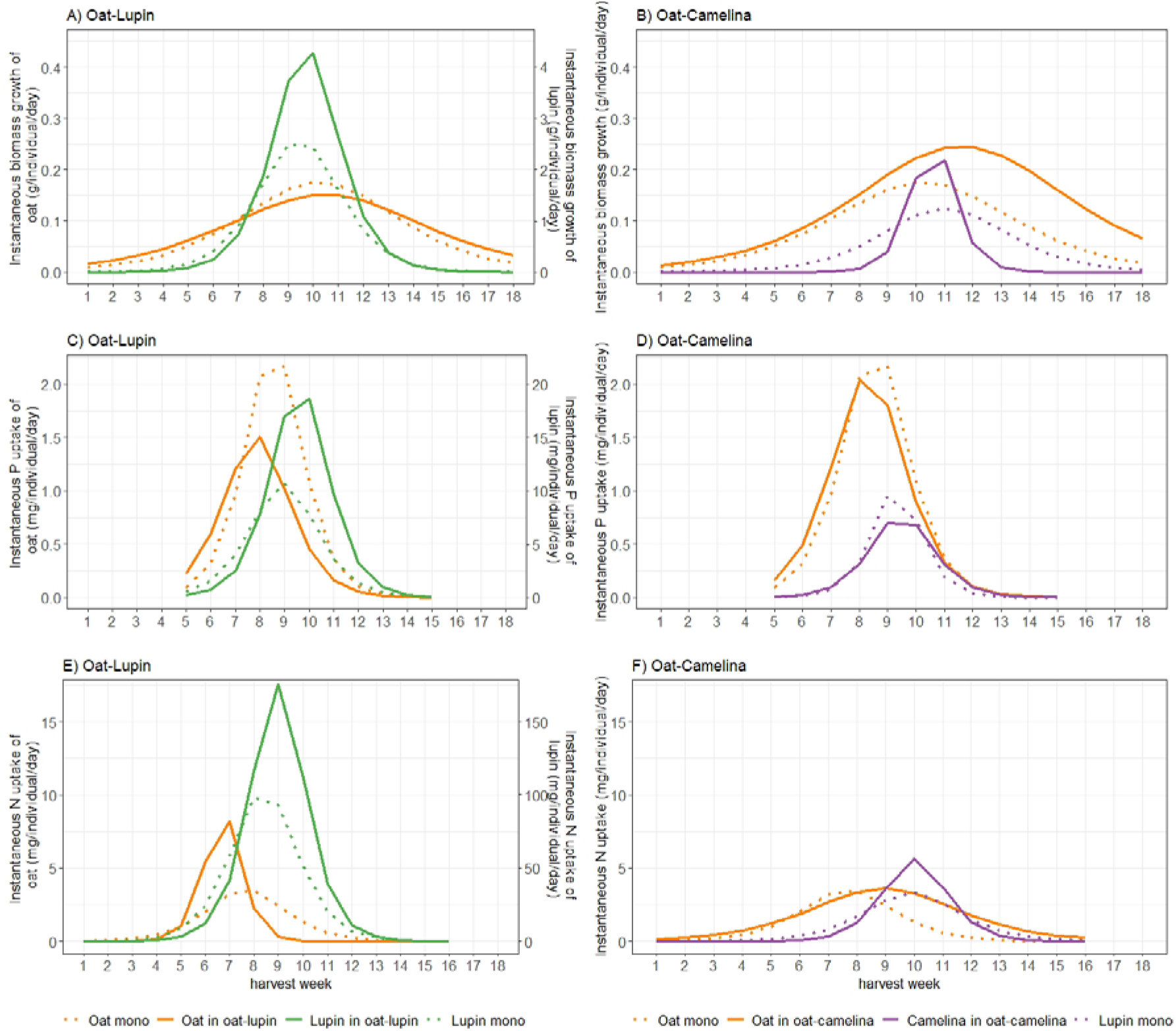
Instantaneous biomass accumulation (A, B), P (C, D) and N (E, F) uptake of either species in the oat–lupin (A, C, E) and oat–camelina (B, D, F) mixtures (solid lines) and monocultures (dotted lines). Instantaneous biomass growth and N uptake of lupin (A, E) are given on a second x-axis and are a factor 10 higher compared to values of oat.

#### Lupin

Cumulative nutrient uptake and biomass accumulation of lupin mixed with oat and lupin in monoculture were similar until week 9 and afterwards the mixture outperformed the monoculture. Lupin intercropped with oat accumulated 34% more biomass (Fig. 1B), 58% more P (Fig. 1E) and 40% more N (Fig. 1H) than lupin grown in monoculture (Table 1).

Maximum instantaneous biomass accumulation of lupin intercropped with oat was 68% higher than of lupin in monoculture. Similarly, maximum instantaneous uptake of P and N were 84% and 69% higher in intercropped than sole cropped lupin, respectively (Fig. 2A, C, E, Table 1, S3). No temporal differentiation of maximum instantaneous biomass accumulation or nutrient uptake rates between lupin in monoculture and mixture could be observed. Nevertheless, maximum instantaneous nutrient uptake rates and biomass accumulation of isolated single lupin were always more than 10 times higher and were also 1-2 weeks later than when grown in a community (Table 1, S3).

#### Camelina

Camelina accumulated 40% more biomass in monoculture than intercropped with oat (Fig. 1C, Table 1). While no differences could be observed for cumulative P uptake (Fig. 1F), camelina tended to accumulate more N when intercropped with oat than in monoculture (Fig. 1I), although the differences were not statistically significant (Table 1).

Maximum instantaneous biomass accumulation and N uptake of camelina mixed with oat was 102% and 70% higher than camelina in monoculture, respectively (Fig. 2B, F, Table S3). Maximum instantaneous P uptake was 33% higher in monoculture camelina than in intercropped camelina (Fig. 2D, Table S3). No temporal differentiation of maximum instantaneous biomass accumulation or nutrient uptake rates between camelina in monoculture and mixture could be observed (Table 1).

### 3.3. Inter-specific variability

No significant differences in the timepoints of maximum instantaneous nutrient uptake rates in the oat–camelina mixture nor for the timepoints of maximum biomass accumulation rates in both mixtures were observed (Table 1). However, maximum instantaneous N and P uptake rates of oat were ~2 weeks earlier than that of lupin in the oat–lupin mixture.

### 3.4. Nitrogen fixation and root exudation

Biological N_2_-fixation differed significantly among diversity levels and a post-hoc test revealed that biological N_2_-fixation was lower in isolated single lupins compared to lupin grown in mixture or monoculture (Table 2, S4). There were no differences in N_2_ fixation between harvest weeks for lupin grown in mixture, monoculture or isolated single plants (Table S4).

**Table 2:** Mean nitrogen derived from the atmosphere (Ndfa)[%] ± SE (n=3) per harvest week for lupin grown in mixture and lupin grown in monoculture.

| harvest week | Ndfa (%) |  |  |
| --- | --- | --- | --- |
|  | lupin in oat–lupin | lupin in monoculture | lupin as isolated single |
| 5 | 83.06 ± 3.36 | 77.16 ± 2.33 | 44.9 ± 3.72 |
| 6 | 76 ± 3 | 69.33 ± 3.96 | 43.83 ± 14.43 |
| 7 | 79.41 ± 5.52 | 70.63 ± 4.26 | 76.63 ± 13.16 |
| 8 | 78.98 ± 3.46 | 72.67 ± 6.93 | 57.69 ± 1.39 |
| 9 | 76.07 ± 1.13 | 76.91 ± 3.69 | 57.39 ± 12.23 |
| 10 | 76.51 ± 6.75 | 74.13 ± 6.39 | 47.67 ± 12.79 |
| 11 | 72.38 ± 3.56 | 70.27 ± 8.03 | 48.6 ± 10.99 |
| 12 | 67.18 ± 10.99 | 54.03 ± 6.62 | 38.66 ± 7.99 |
| 13 | 74.1 ± 7.1 | 66.87 ± 16.63 | 45.71 ± 7.61 |
| 14 | 76.63 ± 13.16 | 76.53 ± 6.64 | 36.91 ± 12.45 |
| 15 | 72.68 ± 6.57 | 71.21 ± 8.41 | 37.14 ± 11.14 |
| 16 | 64.92 ± 8.17 | 70.05 ± 3.66 | 28.14 ± 5.11 |

Succinate, malate, acetate, lactate and citrate were detected as organic acids. Overall exudation of organic acids during the entire growing season was highest for camelina followed by oat and lowest for lupin (Fig. 3). Although not significant, oat in monoculture tended to exude more organic acids early in the growing season (week 4-7) than when grown in mixture with either lupin or camelina (Fig. 3A). Organic acid root exudation for lupin was similar whether lupin was grown in monoculture or intercropped with oat and did not show significant fluctuations throughout the growing season (Fig. 3B). Camelina intercropped with oat exuded more organic acids and earlier compared to when grown in monoculture (Fig. 3C). Although camelina exuded larger amounts of organic acids than both, lupin and oat, this higher exudation was not accompanied by a significant drop in rhizosphere pH (Fig. S2).

**Fig. 3:**
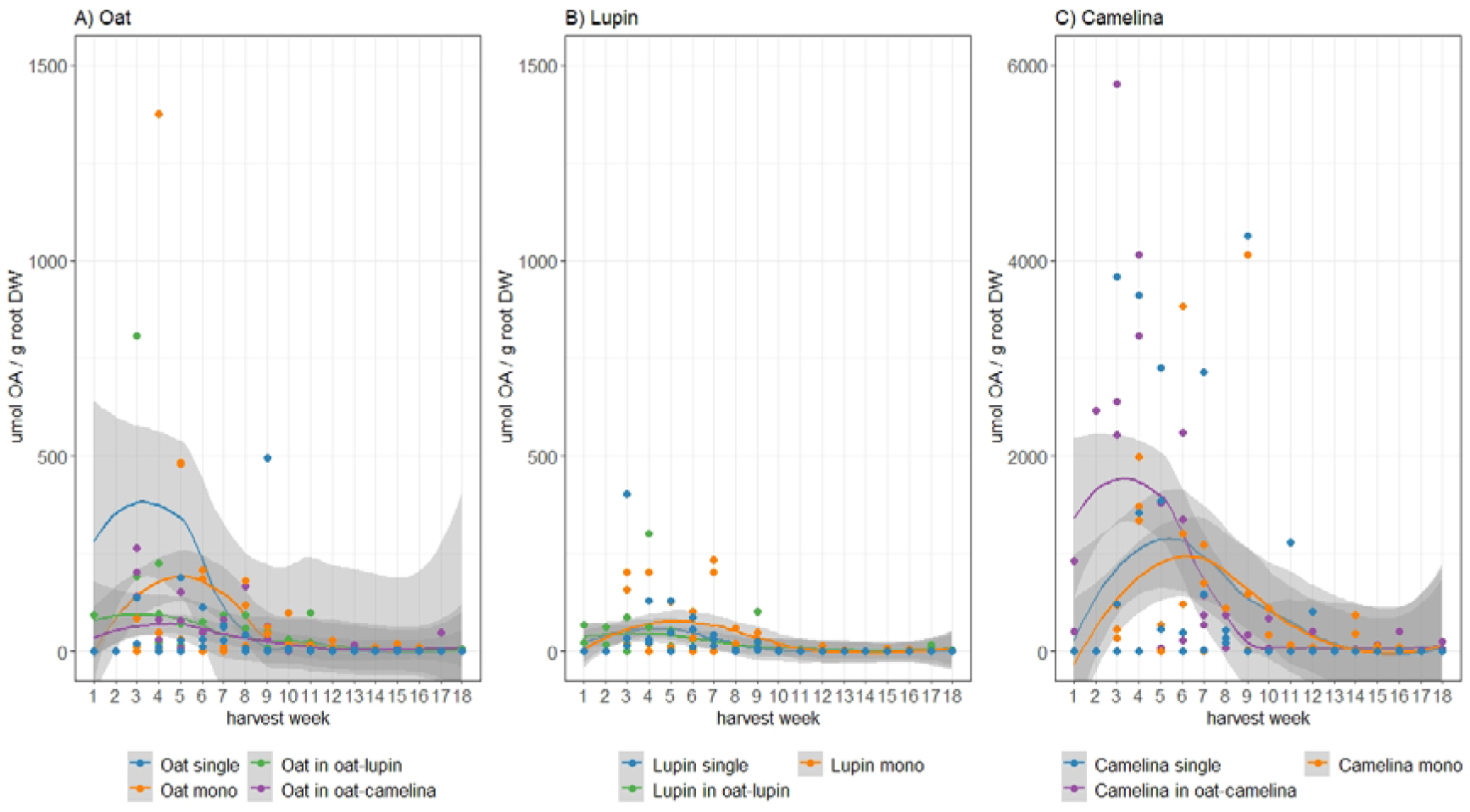
Total organic acids (OA) found in rhizosphere soil in μmol / g root dry weight of oat (A), lupin (B) and camelina (C) grown as isolated single plants (blue), monocultures (orange) and mixtures (green / purple). Note differing y-axis scales. Shading refers to standard errors computed using a t-based approximation.

## 4. Discussion

The methodological approach first applied by Trinder *et al.* (2012) allowed us to analyze the dynamics of competitive plant resource capture and biomass accumulation at two different diversity levels and compare them to isolated single plants that did not experience competitive interactions. We found that competition significantly reduced maximum nutrient and biomass accumulation of plants growing in a community (mixture or monoculture). These plants also showed earlier maximum nutrient uptake and biomass growth, a temporal shift we interpreted as being due to competitive interactions, with uptake and growth trajectories curtailed by competition among and between species.

While both mixtures in this study overyielded, the pathways to overyielding were quite different. Based on partial LERs, oat benefited from intercropping and this benefit was stronger when intercropped with camelina than when intercropped with lupin. Lupin benefited from intercropping with oat but camelina did not. In the oat–lupin mixture, oat initially profited from the slower establishment of the lupin and lupin’s N coming to 70-80% from biological N_2_-fixation. Gradually, however, lupin became a stronger competitor, outcompeting oat for P uptake. Oat N uptake also slowed with time, resulting in a final accumulated oat biomass similar to that in monoculture; while the intercropped lupin strongly overyielded. While the oat–lupin mixture was characterized by strong interactions, the oat–camelina mixture was characterized by temporal partitioning during the early growth stages and aboveground competition in the later growth stages. Here, overyielding of intercropped oat was not due to belowground competitiveness for nutrients, but rather to stronger aboveground competitiveness for light during the later growth stages. Exudation of organic acids did not increase P uptake by oat, although higher exudation by intercropped camelina could have improved N availability via microbial pathways.

### Intra-specific variability

The significantly (up to 30 times) higher maximum accumulated biomass of isolated single plants compared to plants in a community, indicated the scale of the effect of intra-specific competition on nutrient uptake and biomass accumulation in crop systems.

Isolated single lupins accumulated less N from biological N_2_-fixation than lupins in a community. Isolated lupins experienced no competition for soil N, and so might not have invested resources into relatively costly biological N_2_-fixation (Vitousek & Field 1999). However, in contrast to other studies (Corre-Hellou, Fustec & Crozat 2006; Hauggaard-Nielsen *et al.* 2008; Hauggaard-Nielsen *et al.* 2009), we observed no difference in N_2_-fixation between lupin grown in mixture or monoculture, nor any temporal fluctuations, indicating that lupin in a community relied to a consistent level on biological N_2_-fixation.

With respect to temporal shifts, we always found earlier peaks in nutrient uptake and biomass accumulation rates when species were grown in a community. These results contrast with those of Trinder *et al.* (2012) who, in a study using *Dactylis glomerata* and *Plantago lanceolata*, found delay in *D. glomerata* and advancement in *P. lanceolata* of maximum nutrient uptake and biomass accumulation rates when grown in competition. When grown together, the later species, *D. glomerata*, took up more N and suffered less restricted biomass accumulation, indicating it was competitively stronger despite having later peaks in uptake and accumulation rates. In our study, earlier peak rates in a community were always accompanied by significantly lower nutrient and biomass accumulation. Thus, we interpreted the uniform shift towards earlier peak uptake and growth rates for plants in our communities as a passive response to competition, with the onset of competitive interactions between the intercropped species curtailing the trajectories of nutrient uptake and biomass growth, resulting in lower accumulation. This contrasts with the apparently active response found by Trinder *et al.* (2012) where Dactylis shifted uptake and growth to a more favorable timepoint when in competition.

The study of Trinder *et al.* (2012) was conducted with perennial grassland species, and annual crop species might react differently to neighbor presence. For example, a study using two barley cultivars – one early and one late - grown as isolated single plants and in either intra- or inter-specific competition (Schofield *et al.* 2019b) found peak N accumulation was advanced by 0.5 days for the early and delayed by 14.5 days for the late cultivar when in intra-specific competition, while no shifts were observed in inter-specific competition. This suggests crop species may have enough temporal plasticity to avoid competition with kin but not with other species. However, studies contrasting temporal dynamics between isolated and competing plants are extremely rare, and so generalities are difficult and the mechanisms behind these processes remain unknown, indicating further research is needed in this area.

### Oat–camelina mixture

Oat in monoculture yielded 38% less than in mixture with camelina, despite accumulating similar amounts of P, suggesting P was not a comparatively limiting nutrient in monoculture. However, accumulated N of intercropped oat was significantly higher than monoculture oat. Intercropped camelina accumulated less biomass than camelina in monoculture, but this was not due to nutrient accumulation, which did not differ between camelina in monoculture and mixture (Table 1). Thus, we can exclude the idea of oat being a stronger competitor for soil N, which should have resulted in lower N uptake by the intercropped camelina. Instead, the absence of a negative effect of the over-yielding oat on camelina’s nutrient uptake could indicate a partitioning of belowground resources during early growth. While this belowground partitioning could have been spatial (i.e. different rooting depths (Kutschera, Lichenegger & Sobotik 2018)), we found some evidence for temporal partitioning, with intercropped oat accumulating N and P earlier than intercropped camelina, translating into an earlier accumulation of biomass by the oat (Fig. 2). Earlier accumulation of biomass by the oat could then have resulted in shading of the intercropped camelina, reducing camelina growth later in the growing season. Thus, belowground temporal niche separation could have developed into aboveground competition during later growth stages. Such a shift from a positive, belowground effect early in the season to a negative, late-season aboveground effect has been observed in a study of the interactions between barley and the rare arable weed *Valerianella rimosa* (Brooker *et al.* 2018): early in the season barley had a positive, soil-driven effect on *V. rimosa* abundance but with time this shifted to growth suppression by barley, most likely due to light competition (Brooker *et al.* 2018).

Besides belowground partitioning, exudation of organic acids could have played an additional role in the oat–camelina mixture. Early in the growing season, intercropped camelina exuded almost twice as many organic acids as camelina in monoculture, perhaps improving N availability for both species. In a follow-up study to their (2019b) experiment, Schofield *et al.* (2019a) also examined temporal dynamics of soil microbial enzyme activity in the same system and found that temporal dynamics of plant resource capture and soil microbial activity were linked. However, while root exudates can influence soil N availability via microbial pathways (Meier, Finzi & Phillips 2017), root-microbe-soil nutrient interactions are complex, and our understanding of the exact mechanisms limited (Zhang, Vivanco & Shen 2017).

### Oat–lupin mixture

The observed yield advantage for oat in the oat–lupin mixture agrees with often-observed complementarity in resource use or facilitation in cereal–legume mixtures (Duchene, Vian & Celette 2017). We observed that oat intercropped with lupin initially accumulated N at a faster rate than oat grown in other combinations, which is explained by the slow establishment of lupin (Fig. 1) and the general observation that biological N_2_-fixation in lupin only starts four to five weeks after emergence, after which N does not accumulate in shoots for two further weeks (Walker *et al.* 2011). Perhaps because biological N_2_-fixation is a P-demanding process (Walker *et al.* 2011), we observed a close link between the instantaneous N and P uptake rates of intercropped lupin (Fig. 2) and a strong competitiveness of intercropped lupin compared to oat for soil P, resulting in lower accumulated P in lupin–intercropped oat (Fig. 1). However, despite reductions in P uptake, the biomass of lupin– intercropped oat was not negatively affected, and it still accumulated more N than in monoculture. We suggest this latter effect was due to lupin capturing ~70-80% of its N from biological N_2_-fixation.

Surprisingly, N accumulation of oat intercropped with lupin stopped after week nine, even though lupin captured N from biological N_2_-fixation throughout the experiment. There are two possible explanations. First, intra-specific competition among oats could have limited N accumulation at that point in time. Intra-specific competition for N is also visible in the oat monoculture and, although occurring slightly later than in the lupin-intercropped oats, it results in lower final N accumulation, an effect which may be explained by intercropped oats having the advantage of growing with a slowly-establishing legume. The second explanation is that after slow establishment, lupin rapidly accumulated biomass which shaded the oats and reduced N uptake. As N accumulation is closely related to photosynthetic capacity (Sinclair & Horie 1989), reduced N uptake may reflect limited photosynthetic capacity of oats due to light competition with lupin. However, from our data we cannot differentiate between the two.

## 5. Conclusion

This study shows how temporal dynamics of resource uptake and biomass accumulation throughout the growing season can shed light on competitive plant–plant interactions and improve our understanding of the underlying processes that drive yield advantages in intercropping systems. We showed that trajectories of nutrient uptake and biomass accumulation were curtailed by competition, leading to earlier peak rates and lower total accumulated nutrients and biomass. Our results also revealed multiple pathways to overyielding. We observed strong competitive interactions in the oat– lupin mixture, but the oat–camelina mixture was characterized by an apparent shift from positive, belowground temporal partitioning early in the growing season to aboveground competitive interactions later in the growing season. While this study focused on the temporal dynamics of nutrient uptake, biomass accumulation, biological N_2_-fixation and root exudation, our results suggest that temporal patterns of aboveground competitive interactions for light could be equally important for understanding plant–plant interactions in intercropping systems. Understanding temporal dynamics of below- and aboveground resource uptake and biomass accumulation in intercropped plant communities will enable us to maximize the benefits from intercropping systems, as it will facilitate the optimization of these systems with respect to nutrient inputs, enabling us to better design efficient species combinations where temporal differentiation can reduce competition between species and thus increase yields.

## Acknowledgements

We are grateful to Sandra González Sánchez, Jianguo Chen and Zita Sartori for their help with the field experiment, and Anna Bugmann for her help with the lab work. We also thank Asociación Aprisco, the ETH research facility in Lindau and the Sustainable Agroecosystems group for allowing us to use their land and greenhouse facilities. The study was funded by the Swiss National Science Foundation (PP00P3_170645).

## Author contributions

N.E. and C.S. conceived the study with input from L.S. and R.W.B. N.E and L.S. collected the data. N.E. and B.S. analyzed samples in the laboratory. N.E. assembled and analyzed the data with the help of C.S and R.W.B. N.E. wrote the first draft of the paper. All authors discussed data analyses and results and revised the manuscript.

