## Supplement for "Temporal differentiation of resource capture and biomass accumulation as a driver of yield increase in intercropping"

**Supplementary material**

**Tables**

*Table S1: Parameters indicating the fit of logistic nls models. AvLu: oat with lupin, AvCa: oat with camelina, AvAv: oat monoculture, LuAv: lupin with oat, LuLu: lupin monoculture, CaAv: camelina with oat, CaCa: camelina monoculture. RSS: Residual sum of squares, R^2^: ratio of residual sum of squares (RSS) to total sum of squares (TSS). #it: Number of iterations to reach model convergence. Df: degrees of freedom.*

| **Biomass** | | | | | | |
| --- | --- | --- | --- | --- | --- | --- |
|  | RSS | R^2^ | # it. | Achieved convergence tolerance | Residual Std. Error | Df |
| AvLu | 5.84 | 0.74 | 0 | 8.36E-06 | 0.3384 | 51 |
| AvCa | 7.79 | 0.82 | 0 | 4.26E-06 | 0.3948 | 50 |
| LuAv | 1026.36 | 0.7 | 0 | 5.69E-06 | 4.486 | 51 |
| CaAv | 6.29 | 0.29 | 6 | 8.44E-06 | 0.3548 | 50 |
| AvAv | 3.79 | 0.81 | 0 | 8.78E-06 | 0.2724 | 51 |
| LuLu | 324.18 | 0.79 | 4 | 8.94E-06 | 2.521 | 51 |
| CaCa | 3.88 | 0.54 | 0 | 4.58E-06 | 0.2786 | 50 |
| Av | 1860.5 | 0.59 | 1 | 4.84E-06 | 6.1 | 50 |
| Lu | 43182.8 | 0.8 | 2 | 6.99E-06 | 29.1 | 51 |
| Ca | 4949.76 | 0.43 | 5 | 8.46E-06 | 9.852 | 51 |
| **Phosphorous** | | | | | | |
|  | RSS | R^2^ | # it. | Achieved convergence tolerance | Residual Std. Error | Df |
| AvLu | 207.02 | 0.33 | 4 | 3.62E-06 | 2.627 | 30 |
| AvCa | 89.93 | 0.72 | 1 | 3.76E-06 | 1.731 | 30 |
| LuAv | 7064.28 | 0.72 | 14 | 9.58E-06 | 16.18 | 27 |
| CaAv | 75.82 | 0.24 | 9 | 8.7E-06 | 1.646 | 28 |
| AvAv | 280.63 | 0.45 | 4 | 7.92E-06 | 3.111 | 29 |
| LuLu | 3339.77 | 0.68 | 3 | 6.68E-06 | 10.55 | 30 |
| CaCa | 56.19 | 0.34 | 7 | 8.12E-06 | 1.369 | 30 |
| Av | 9409.82 | 0.39 | 0 | 5.3E-06 | 17.71 | 30 |
| Lu | 230553.4 | 0.83 | 4 | 7.36E-06 | 92.41 | 27 |
| Ca | 19746.9 | 0.2 | 0 | 5.6E-06 | 31.42 | 20 |
| **Nitrogen** | | | | | | |
|  | RSS | R^2^ | # it. | Achieved convergence tolerance | Residual Std. Error | Df |
| AvLu | 2879.2 | 0.47 | 3 | 6.3E-06 | 8.089 | 44 |
| AvCa | 1656.65 | 0.71 | 5 | 6.75E-06 | 6.136 | 44 |
| LuAv | 1394605 | 0.64 | 5 | 7.12E-06 | 176 | 45 |
| CaAv | 6349.42 | 0.24 | 14 | 8.37E-06 | 12.44 | 41 |
| AvAv | 686.38 | 0.72 | 1 | 8.79E-06 | 3.905 | 45 |
| LuLu | 444691.9 | 0.73 | 10 | 8.83E-06 | 99.41 | 45 |
| CaCa | 1005.7 | 0.6 | 15 | 9.54E-06 | 4.953 | 41 |
| Av | 234828.9 | 0.62 | 1 | 4.57E-06 | 77.6 | 39 |
| Lu | 83597206 | 0.69 | 3 | 9.57E-06 | 1446 | 40 |
| Ca | 1125044 | 0.41 | 1 | 7.74E-06 | 197 | 29 |

*Table S2:Mean δ ^15^N values ± SEM of leaves of lupin that are fully dependent upon N_2_ fixation and sampled at the same harvest week as the field plants (β-values). n = 3 per week.*

| harvest week | *β* |
| --- | --- |
| 1 | 1.33 ± 0.06 |
| 2 | 0.8 ± 0.26 |
| 3 | 0.82 ± 0.26 |
| 4 | -0.22 ± 0.04 |
| 5 | -0.43 ± 0.1 |
| 6 | -0.79 ± 0.39 |
| 7 | -0.54 ± 0.08 |
| 8 | -0.71 ± 0.15 |
| 9 | -0.84 ± 0.1 |
| 10 | -0.84 ± 0.11 |
| 11 | -0.94 ± 0.06 |
| 12 | -0.94 ± 0.1 |
| 13 | -1.1 ± 0.24 |
| 14 | -1.23 ± 0.1 |
| 15 | -1.38 ± 0.22 |
| 16 | -1.55 ± 0.15 |

*Table S3: I_max_* *values for each species in mixture, monoculture or as isolated single plant. I_max_ is the maximum instantaneous biomass accumulation (g day^-1^) and nutrient uptake rate (mg day^-1^) which emerges at the time t_max_.*

|  | Biomass | Phosphorous | Nitrogen |
| --- | --- | --- | --- |
| Oat : Lupin | 0.15 | 1.51 | 8.89 |
| Oat : Camelina | 0.25 | 2.11 | 3.61 |
| Oat mono | 0.18 | 2.35 | 3.56 |
| Oat single | 2.91 | 6.59 | 42.67 |
| Lupin : Oat | 4.38 | 19.78 | 175.76 |
| Lupin mono | 2.61 | 10.73 | 103.85 |
| Lupin single | 69.04 | 284.7 | 2684.4 |
| Camelina : Oat | 0.25 | 0.77 | 5.65 |
| Camelina mono | 0.13 | 1.03 | 3.33 |
| Camelina single | 3.99 | 34.99 | 116.62 |

*Table S4: Results of linear mixed effects ANOVA testing the effects of diversity (mixture, monoculture, isolated singles) and harvest week on %Ndfa. Random term was replicate. SS: Sum of squares, MS: mean of squares, Df: degrees of freedom, F-value: variance ratio. P-values in bold are significant at α = 0.05 (* P < 0.05, ** P < 0.01, *** P < 0.001).*

|  | Df | SS | MS | F-value | P |
| --- | --- | --- | --- | --- | --- |
| ***diversity*** | 2 | 15591.9 | 7796 | 47.267 | **<0.001***** |
| *harvest week* | 11 | 3032.7 | 275.7 | 1.672 | 0.1 |
| *diversity × harvest week* | 22 | 2198.4 | 99.9 | 0.606 | 0.905 |

**Figures**

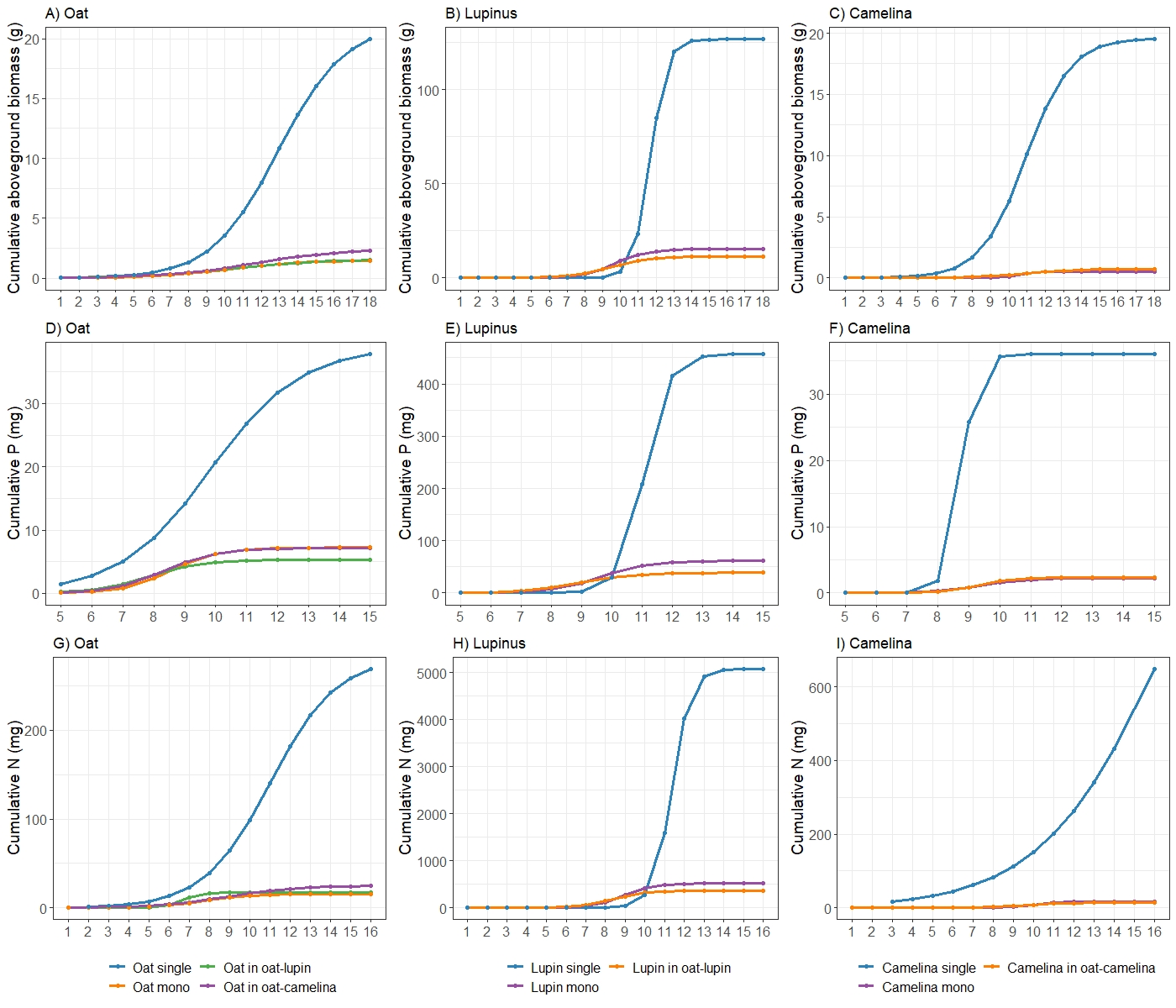

*Fig. S1: Trajectories of cumulative aboveground biomass (A-C), P (D-F) and N (G-I) uptake of oat with lupin (A, D, G: green), oat with camelina (A, D, G: purple), lupin (B, E, H) and camelina (C, F, I) when grown in mixture (purple, green), in monoculture (orange) and as isolated singles (blule). Same data as in Fig. 2 but with addition of isolated singles, which were removed from Fig.2 to aid comparisons between the other treatments. Curves are derived from Eqn. 1. Note different y-axis scales.*

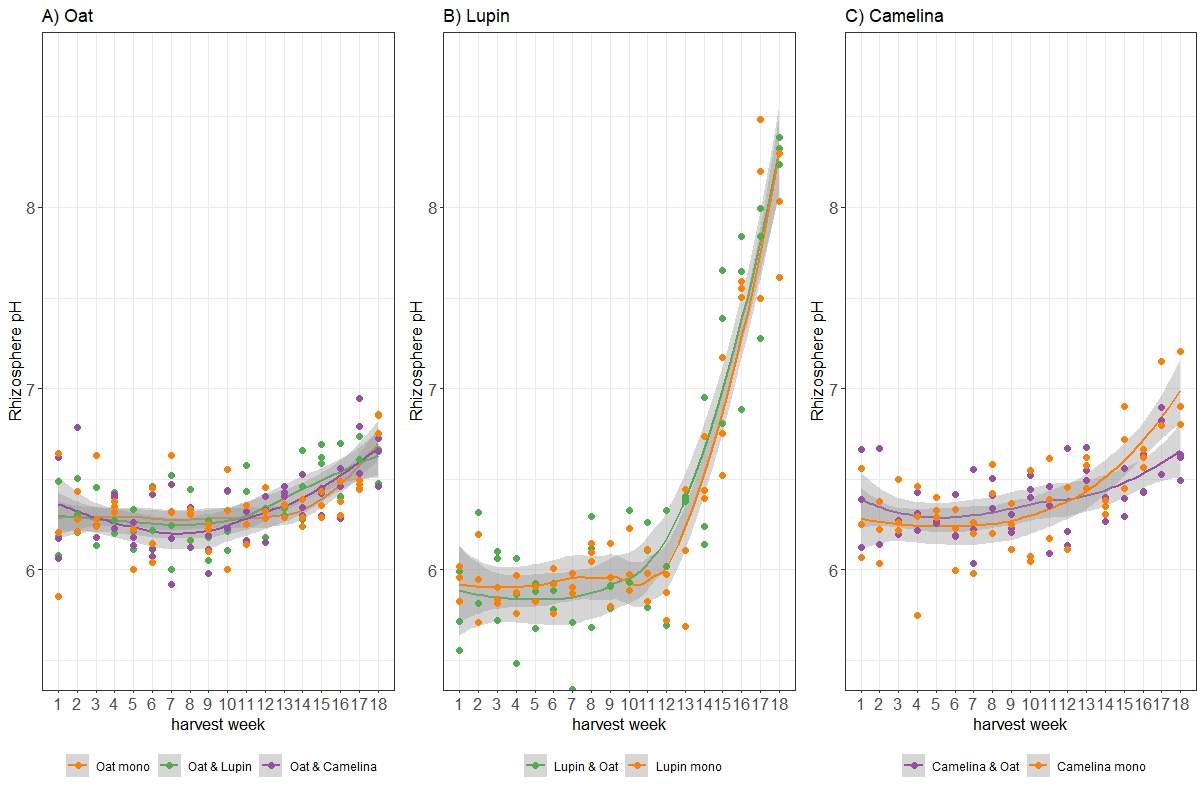

*Fig. S2: Rhizosphere pH measured in rhizosphere soil immersed in 0.2 mM CaCl2 solution for oat (A), lupin (B) and camelina (C) in monoculture (orange) or mixture (green, purple). Shading refers to standard errors computed using a t-based approximation.*
